# Scale-invariant time registration of 24-hour accelerometric rest-activity profiles and its application to human chronotypes

**DOI:** 10.1101/2020.10.13.337550

**Authors:** Erin I. McDonnell, Vadim Zipunnikov, Jennifer A. Schrack, Jeff Goldsmith, Julia Wrobel

**Author notes:** **Corresponding author:** Erin I. McDonnell, Department of Biostatistics, 722 West 168th Street, 6^th^ Floor, New York, NY 10032.

## Abstract

By collecting data continuously over 24 hours, accelerometers and other wearable devices can provide novel insights into circadian rhythms and their relationship to human health. Existing approaches for analyzing diurnal patterns using these data, including the cosinor model and functional principal components analysis, have revealed and quantified population-level diurnal patterns, but considerable subject-level variability remained uncaptured in features such as wake/sleep times and activity intensity. This remaining informative variability could provide a better understanding of chronotypes, or behavioral manifestations of one’s underlying 24-hour rhythm. Curve registration, or alignment, is a technique in functional data analysis that separates “vertical” variability in activity intensity from “horizontal” variability in time-dependent markers like wake and sleep times; this data-driven approach is well-suited to studying chronotypes using accelerometer data. We develop a parametric registration framework for 24-hour accelerometric rest-activity profiles represented as dichotomized into epoch-level states of activity or rest. Specifically, we estimate subject-specific piecewise linear time-warping functions parametrized with a small set of parameters. We apply this method to data from the Baltimore Longitudinal Study of Aging and illustrate how estimated parameters give a more flexible quantification of chronotypes compared to traditional approaches.

## 1. INTRODUCTION

A person’s circadian rhythm is driven by the internal clock that dictates his or her daily physiological cycles, including sleep-wake times, hormone release, core body temperature, blood sugar regulation, and other components of the homeostatic system, all of which influence how one distributes his or her daily activity (Zee et al., 2013). Although circadian rhythms can be influenced by social and solar clocks, they are endogenous and determined largely genetically (Roenneberg et al., 2003). Chronotypes, or phenotypes of circadian rhythms that can be used to identify population subgroups and classify individuals, have been linked to numerous health outcomes. For example, some individuals wake earlier in the day (i.e. a “morningness chronotype”) while others stay awake later into the night (i.e. a “eveningness chronotype”). Those with an eveningness chronotype also tend to be at greater risk for some mental health disorders and general health problems such as hypertension, asthma, and type 2 diabetes (Partonen, 2015).

Chronotypes have typically been defined on a single continuum ranging from morningness to eveningness (Partonen, 2015), but there may be more nuanced phenotypes characterizing one’s pattern of energy over the course of a day, independent of sleep and wake time preferences. For example, a recent study revealed that measures of sleep quality, sleep duration, and diurnal inactivity (e.g. nap times or wakeful rest) showed very little correlation with self-reported chronotype on a scale from “definitely a ‘morning’ person” to “definitely an ‘evening’ person”, suggesting that these other sleep features provide new information beyond the established chronotypes (Jones et al., 2019). The sleep and diurnal features examined were extracted from accelerometers, which give rich data for studying daily activity patterns by collecting epoch-by-epoch (often minute-by-minute) data on activity intensity over 24 hours. Accelerometers are a natural mechanism for tracking diurnal patterns in order to detect underlying circadian rhythms. Novel data-driven techniques are now capable to leverage the rich information collected by accelerometers to confirm the existence of morningness and eveningness chronotypes, to identify more subtle differences in chronotypes, and to illustrate how these are manifested in various epidemiological cohorts and clinical sub-groups.

We introduce a new accelerometer data-driven approach, based on curve registration (Wrobel et al., 2019), for understanding circadian rhythms. Curve registration uses tools from functional data analysis, a statistical subfield that focuses on the kind of intensive longitudinal data made possible by wearable devices and other technologies (Ramsay & Silverman, 2002, 2005). Our method separates and analyzes both 1) “horizontal” variability in time-dependent markers like wake and sleep times, and 2) “vertical” variability in activity intensity; by doing so, the method gives distinct insights into underlying circadian rhythms. The remainder of this paper is organized as follows. In Section 2, we provide an overview of the existing methods that use accelerometer data to study circadian rhythms. In Section 3, we describe the registration method. In Section 4, we use the registration method to analyze accelerometer data from the Baltimore Longitudinal Study of Aging (BLSA). In Section 5, we compare registration to a set of existing methods. We conclude with a discussion in Section 6.

## 2. LITERATURE REVIEW

Circadian rhythms can be studied using a range of data collection instruments, including surveys such as the Munich Chronotype Questionnaire, and measures of core body temperature and hormone levels (Melo et al., 2017; Partonen, 2015; Roenneberg et al., 2003). We highlight methods that involve the collection of minute-by-minute accelerometer data, categorizing them into two distinct classes: landmark methods, which extract predefined features such as sleep and wake times, and methods that use the full 24-hour activity profile in an attempt to quantify the complexity of diurnal patterns.

Landmark methods define chronotypes based on one or two summary measures of a subset of one’s 24-hour period. Some of these methods identify a fixed-length interval, including the 5 least active consecutive hours and the 10 most active consecutive hours, often abbreviated L5 and M10, respectively, in the chronotype literature. The timing of such intervals can indicate morningness versus eveningness preferences (Witting et al., 1990). Other landmark methods identify intervals whose length varies across subjects, such as sleep or daytime intervals (Gershon et al., 2018; Kaufmann et al., 2018; Urbanek et al., 2018). One example is the sleep period time window (SPT-window) approach of van Hees et al. (2015, 2018). The SPT-window, developed for 3-axial raw accelerometer data, is a heuristic approach that entails searching within the 12-hour period centered around the L5 midpoint for periods of “sustained inactivity,” defined by absence of change in arm angle. The SPT-window then stretches from the start of the first period of sustained inactivity to the end of the last period within that 12-hour span. For sleep and daytime intervals, both the duration and the timing of the interval’s midpoint can be assessed, although the midpoint alone is often used to define chronotypes on either a continuous scale (Urbanek et al., 2018) or dichotomously by establishing some cut-off for the midpoint (Gershon et al., 2018; Kaufmann et al., 2018).

All landmark methods suffer from the same drawback: they condense activity data to produce a single, pre-defined aggregate measure (or a few such measures) within long time windows. This can eliminate subtle, but important and relevant differences in daily accelerometry profiles which may help to differentiate, for example, someone who is an early riser but is sedentary in the afternoon from someone who is an early riser and remains continuously active throughout the day. Moreover, only the data inside the landmark interval of interest is summarized, rather than the full 24-hour rest-activity profile. Lastly, for those methods that use only the timing of an interval’s midpoint to define chronotypes, duration of activity and sleep is not factored into the chronotype at all; someone who sleeps from 10PM to 6 AM has the same sleep interval midpoint as someone who sleeps from 11PM to 5AM. Therefore, without separate adjustment for sleep duration, landmark methods may incorrectly classify a broad range of circadian rhythm patterns into a single chronotype.

Other methods utilize 24-hour rest-activity profile, leveraging the complex information and extracting more intricate features of circadian rhythms. The cosinor method captures both the timing and intensity of one’s daily active period in a single three-parameter model. It does so using a parametric regression model with clinically interpretable parameters, including the MESOR (midline estimating statistic of rhythm) or model-based mean intensity, the amplitude or peak intensity, and the acrophase or timing of the peak intensity (Halberg et al., 1967; Marler et al., 2006). The cosinor method has been widely used to study circadian rhythms (Cornelissen, 2014; Refinetti et al., 2007). However, a major limitation of this method is that it assumes all activity curves conform to a cosine shape: this fit may be too restrictive to accurately represent the circadian rhythm profile for many subjects.

Like the cosinor approach, functional principal components analysis (FPCA) models epoch-by-epoch accelerometry data; however, FPCA is a data-driven approach that does not impose any pre-defined parametric assumptions about the shape of daily rest-activity rhythms. Each person’s 24-hour rest-activity profile is represented via a combination of common population-level patterns (principal components) that capture most of the variability across subjects representing the population. Principal component (PC) scores for a given subject each indicate the degree to which these patterns are present for that subject, and tell a piece of the circadian rhythm puzzle: whether he or she has a generally more active day, whether he or she wakes up early or late, how long his or her day is, and other specific and independent aspects of the circadian rhythm. FPCA has recently gained traction within the sleep research community (Gershon et al., 2016; Zeitzer et al., 2013), in part due to the appeal of its flexibility. However, 24-hour rest-activity profiles have two distinct sources of variability: horizontal variability, which reflects the timing of features like wake and sleep times and peaks of activity, and vertical variability, which reflects the intensity of activity. FPCA does not separate these distinct sources of variability, and the patterns that are identified can conflate the two sources in ways that could mask the true behavioral differences between people.

The registration method introduced in this paper retains the strengths of existing approaches while addressing their limitations. By using the full 24-hour rest-activity profile, registration can identify differences in landmarks such as sleep and wake times, but also provides more detailed information than landmark methods. In contrast to the cosinor approach, registration does not place a restrictive shape on the data, but still produces interpretable and clinically meaningful parameters. Registration builds on FPCA as a standalone method by separating horizontal variability (timing of activity) from vertical variability (intensity of activity), which results in more easily interpretable PCs. We will show that the registration method, by explicitly separating the sources of variability, reveals new insights into accelerometry-estimated chronotypes and circadian rhythms.

## 3. THE REGISTRATION METHOD

### 3.1 Overview

Our approach brings two new ideas to the study of circadian rhythms. First, we dichotomize the minute-level activity counts rather than directly analyzing minute-level (or other epoch-level) activity counts like the methods discussed in the previous section. When activity counts are analyzed directly, the periods with high-intensity exercise or other strenuous activities may heavily influence the shape of one’s activity profile and consequently our inference about one’s chronotype. Another limitation of existing approaches that they provide results that depend on the definition (scale) of activity count that is often device- and manufacturer-specific. Our approach overcomes this limitation by dichotomizing minute-level activity-counts into “active” or “resting” states with the objective of detecting patterns in the probability of active state over time, which we term “activity probability profiles”. Thus, our approach is scale-invariant. Note that we use “resting” to refer both to “sedentary” (awake, but not active) and sleep periods.

Next, we introduce registration in the context of accelerometry-estimated circadian rhythms. The goal of our method is to align subjects’ activity probability profiles – smooth, underlying curves that quantify the probability that a subject is active at each time of the day – based on shared features such as wake and sleep times and peaks in activity probability. We accomplish this by separating the variability across activity probability profiles into the horizonal and vertical sources. Figures 1 and 2 illustrate this process. In the left panel of Figure 1, each curve represents a different subject’s activity probability profile over a 24-hour period, with one subject highlighted in red. Differences across subjects in the timing of common features such as sleep and wake times make it difficult to identify any shared daily patterns. In the right panel of Figure 1, we have used the proposed registration approach to align these curves and a clearer shared pattern is revealed, showing a morning peak, a midday dip in activity, and an afternoon peak; nearly all variability in the right panel of Figure 1 is vertical variability. The transformation of curves in the left panel to curves in the right panel is accomplished through subject-specific (curve-specific) timewarping functions shown in the middle panel of Figure 1, which capture the horizontal variability.

**Figure 1.**
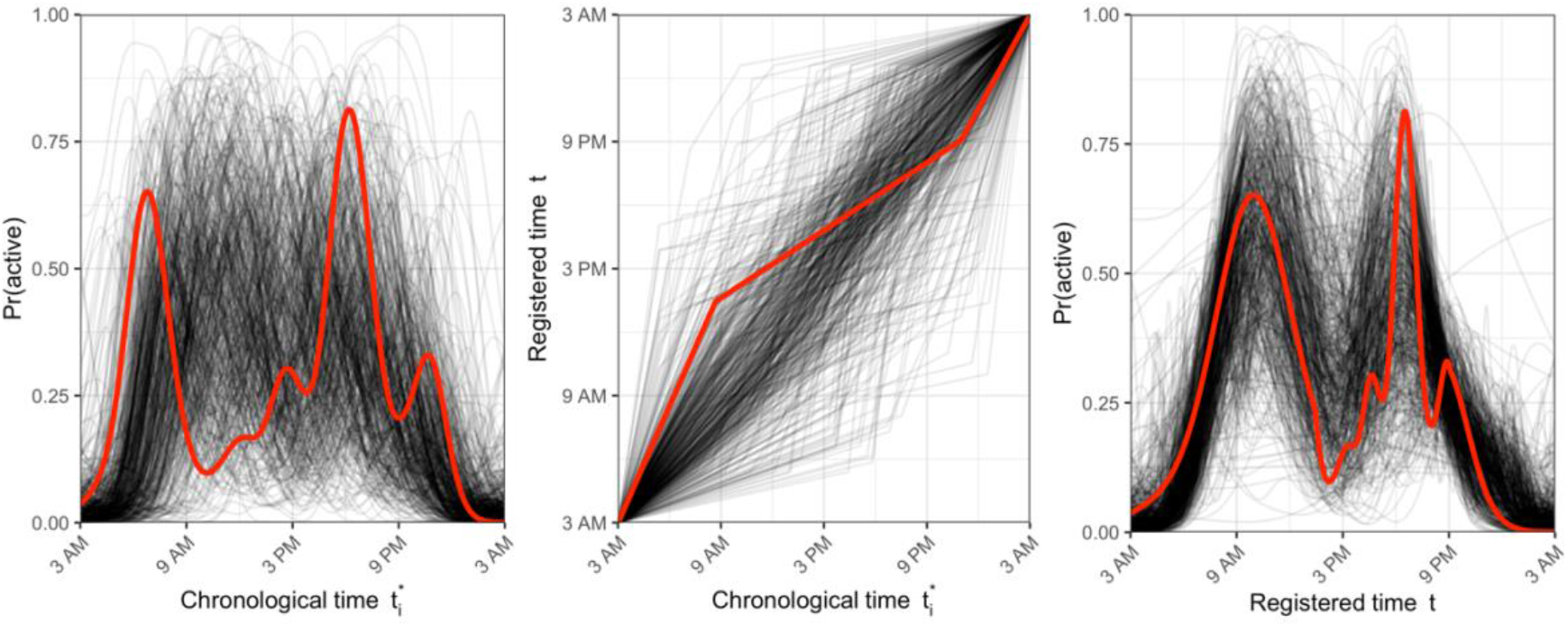
Activity probability profiles pre-registration (left), warping functions (middle), and activity probability profiles post-registration (right). In each panel, each curve represents data for a particular subject. Data from one subject is highlighted in red across all panels.

Warping functions map observed “chronological” time to a latent “registered” time domain on which all profiles are aligned. Figure 2 illustrates the effect of the warping function for the subject highlighted in Figure 1. This subject’s window of activity between 3AM and 9AM in chronological time is stretched, or slowed down, by the warping function, so that it covers roughly 3AM through 2PM in registered time. In other words, the activity profile that this subject experienced in the first 6 hours of their morning, most people take until 2PM to complete. Conversely, the 15-hour window from 9AM to nearly midnight in chronological time becomes compressed or sped up, so that it only accounts for about 8 hours of this person’s active day in registered time. After midnight time goes unchanged, suggesting this subject has an average bedtime relative to the study population. The shape of the warping function in the right panel of Figure 2 reflects this exact mapping from chronological time (x-axis) to registered time (y-axis). Note that our registration framework assumes two time points (knots) that split the 24-hour period into three different periods with different linear dependences (slopes or angles of the linear lines) between the chronological and registered times. Each subject’s set of parameters, i.e. the two knots and three slopes, describe how the timing of his or her activity profile features differ from the population mean. We expand on this in more technical detail below.

**Figure 2.**
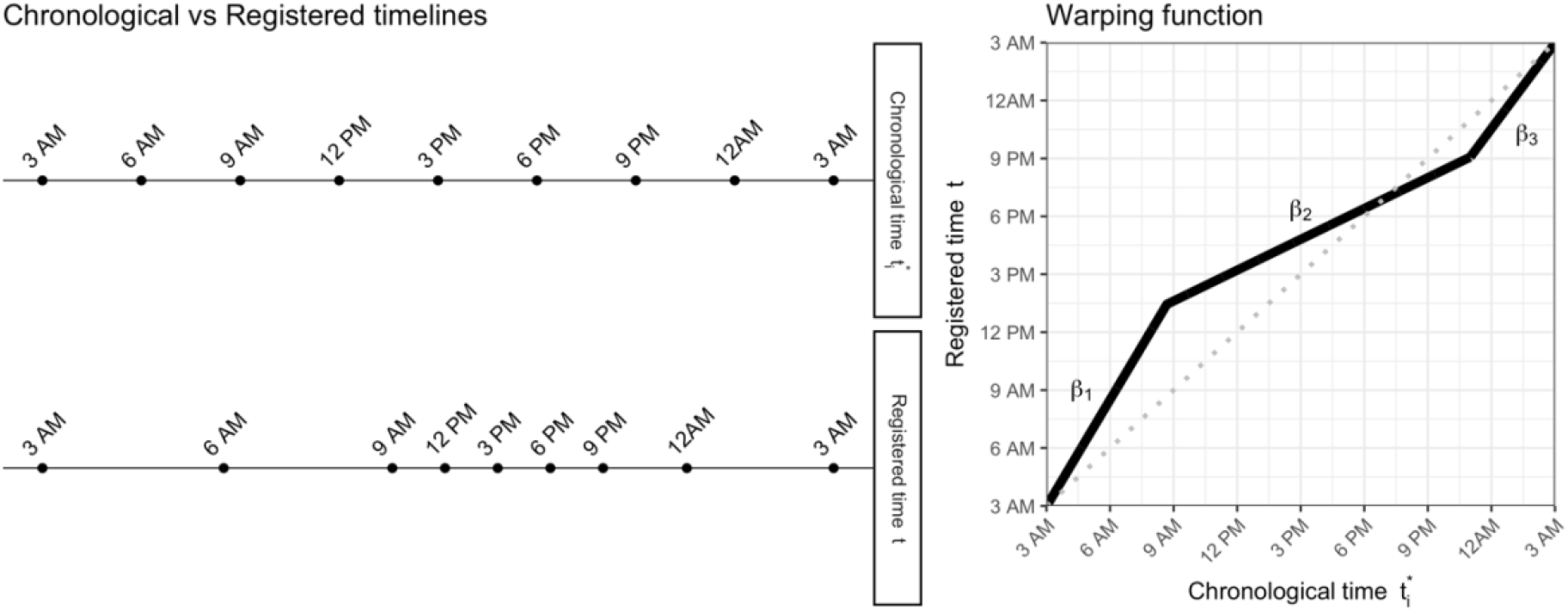
Example of a warping function for one subject mapping chronological time to registered time. The left panel demonstrates where certain points throughout his chronological day fall on his registered timeline. The right panel displays the corresponding 2-knot piecewise linear warping function.

### 3.2 Technical details

Registration accomplishes the separation of horizontal and vertical variability using an iterative two-step approach, cycling between estimating warping functions to capture the horizontal source of variability, and applying FPCA to understand vertical variability in the post-warped/registered patterns. We will begin with the mathematical representation of this two-step iterative process and then provide a more heuristic breakdown of each step. The following notation draws from the statistical subfield of functional data analysis, which provides a useful conceptual framework for intensive longitudinal data (e.g. minute-by-minute rest/active states over 24 hours). Let 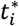 represent chronological time and let 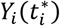 represent the *i^th^* subject’s activity probability profile over chronological time.

Warping functions 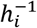 map observed chronological time 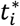 to registered time *t*. The resulting unregistered and registered activity probability profiles can be written as 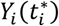 and 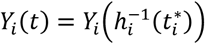, respectively; this notation implies activity probability profiles observed over continuous time, although in practice we observe these profiles at discrete times (for example, at minute level). The warping functions align the observed activity probability profiles to subject-specific mean templates *μ_i_*(*t*). which are estimated using a binary form of FPCA. Notationally, we combine warping functions with FPCA through the following:

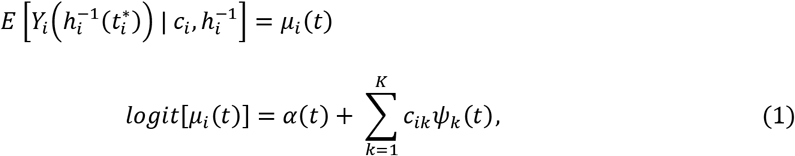

where *ψ_k_*(*t*) are the PC functions, *c_ik_* are subject-specific PC scores, and *α*(*t*) is the population mean function. In the first step of Equation (1), we estimate the warping functions 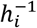 while holding the mean templates *μ_i_*(*t*) fixed. In the second step, we estimate the mean templates *μ_i_*(*t*) using FPCA on the registered data while holding 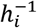 fixed. We iterate through these two steps until convergence. Below we explain these two steps in greater detail; a full technical overview of registration can be found in (Wrobel et al., 2019).

#### 3.2.1. Registration Step 1. Warping functions

In the first step of Equation (1), we estimate subject-specific warping functions 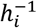 to map chronological time 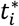 to registered time *t* so that the resulting registered activity probability profiles are aligned to the most recent mean templates *μ_i_*(*t*). The warping functions can take many forms; in this paper, we define a 2-knot piecewise linear function that is novel in the literature, elegant in its simplicity and interpretability, and flexible enough to accurately register daily activity profiles. As demonstrated in Figures 1 and 2, the function for a given subject *i* consists of a series of three slopes {*β*_1*i*_, *β*_2*i*_, *β*_3*i*_} connected at two knot locations {*k*_1*i*_, *k*_2*i*_} along the chronological timeline:

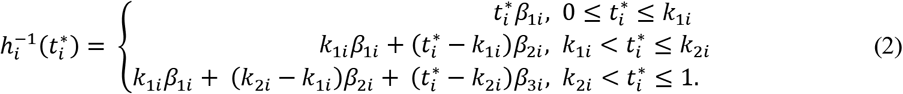

For simplicity, we rescale the 24-hour period so that time is defined on [0,1]. We place additional constraints on the warping functions to ensure that they are always increasing (i.e. we never warp time backwards) and that times 0 (the beginning of the 24-hour period) and 1 (the end of the 24-period) always map to 0 and 1, respectively. The three slopes and two knots that make up each subject’s warping function convey information about his or her diurnal pattern relative to the population.

#### 3.2.2. Registration Step 2. FPCA

In the second step of Equation (1), we use FPCA to estimate subject-specific mean templates *μ_i_*(*t*), using the most recent warping functions to define registered time 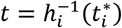. In other words, we perform FPCA on the newly registered activity probability profiles 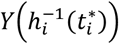. Because we propose to dichotomize the activity counts, we use the extension of FPCA that applies to binary functional data (Goldsmith et al., 2015; van der Linde, 2008). Binary FPCA estimates a functional template *μ_i_*(*t*) of the probability of activity over registered time for each subject, which reflects the population mean behavior while allowing for subject-specific deviation. Specifically, the logit of *μ_i_*(*t*) is decomposed into a population-level functional mean *α*(*t*), *K* population-level functional PCs *ψ*_1_(*t*),…, *ψ_K_*(*t*). and subject-specific scores *c*_*i*1_,…, *c_ik_* which indicate how much each component contributes to the subject-specific activity probability template.

#### 3.2.3. Iterate between the warping function and FPCA steps

Registration iterates between these two steps of Equation (1) until the activity probability curves are temporally aligned across subjects. After registration, we have separated the original activity probability profiles into warping functions and aligned activity probability profiles, which can each be used to understand different aspects of individual variation in circadian rhythms. From the warping functions, subject-specific slopes reveal information about the increasing/decreasing nature of subject-specific time with respected to the registered time and the two times of the day when the linear slopes defining the subject-specific alignment change. From the aligned activity profiles, we can learn about both population-level and subject-specific activity patterns. The estimated population mean activity profile and PC functions describe a typical 24-hour activity probability profile and the major sources of variability across subjects, respectively. The subject-specific PC scores indicate individual deviations from the population mean profile after removing any variability in the timing of landmark features. All of these pieces of information contain useful insights into heterogenous complex nature of chronotypes across subjects. The registration procedure is implemented in the R package “registr” (Wrobel et al., 2020).

## 4. APPLICATION

Our motivating data come from the Baltimore Longitudinal Study of Aging (BLSA), a prospective observational study collecting health, cognitive, and physical performance evaluations of initially healthy participants every 1-4 years for life. All subjects provided written informed consent to participate in the study. Actiheart activity monitors (Schrack et al., 2014) were worn by the participants for up to seven consecutive days. We illustrate our approach by analyzing the first available Tuesday from each subject, chosen to represent typical weekday activity without the after-weekend inertia typically observed on Mondays (Urbanek et al., 2018). Twenty-four-hour activity periods began at 3AM of the specified day and ended at 2:59AM of the following day. Minute-level activity counts were dichotomized into active (activity count > 10) or resting (activity count ≤ 10) state (Schrack et al., 2018, 2019; Wanigatunga et al., 2019). After data processing, 492 subjects were analyzed. The median age was 71 years (interquartile range: 62 to 80), and 54% were male.

We applied the registration method to these 24-hour rest/active binary profiles, using two PCs for the binary FPCA step. The left, middle, and right panels of Figure 1 display unregistered activity probability profiles obtained by smoothing minute-level rest/active binary curves, estimated warping functions, and registered activity probability profiles, respectively. At the population level, the registered curves in the right panel reveal a common trend across subjects, which includes a morning activity peak, a midday dip during which subjects are in a more restful state, and a late afternoon/evening activity peak. The horizontal variability in activity data is captured in the warping functions and remaining vertical variability in activity probability profiles is captured through the FPCA decomposition of the registered data; these are both interpretable in the context of chronobiology.

The warping functions are parameterized by three subject-specific slopes given in Equation (2), each of which reveal different information about morningness and eveningness. A steep first line segment (slope > 1) indicates that a subject’s activity during the late night and morning gets moved forward in time to match the population pattern — that is, his or her day started earlier in chronological time than the average BLSA subject. An extremely steep first line segment suggests an “early bird”, similar to the example subject from Figures 1 and 2. On the contrary, a warping function with a flatter first line segment (slope < 1) indicates that one’s day started later in chronological time. We can apply analogous interpretations to the third line segment of the warping function, which generally takes place during evening and nighttime hours: extremely steep third line segments indicate “night owls”, while flatter third line segments represent subjects who end their active day earlier than most. When a subject has a warping function close to the identity line (i.e. all three slopes near 1), this suggests that his or her activity probability profile in chronological time was in line with the population mean and therefore needed little to no warping to map to the registered time domain.

The registered activity probability profiles are parameterized by the subject-specific scores from the two PCs estimated using binary FPCA. Figure 3 shows the population mean activity probability profile in black plus (blue) or minus (red) one standard deviation of the first (left panel) or second (right panel) PC score. Together these panels show the patterns identified by FPCA for these data. The first PC captures variability across subjects in terms of overall activity probability. The second PC captures variability across subjects in terms of which activity peak (morning or evening) is more dominant. This should not be confused with morningness vs. eveningness chronotypes, which are a reflection of sleep timing (and are better captured by warping function parameters). Some remaining variability in wake times that was not captured in the warping functions is indeed evident in this second PC, as well as some potential restlessness at the end of the activity profile. However, the second PC largely indicates whether a subject is most consistently active immediately upon waking or later in the day.

**Figure 3.**
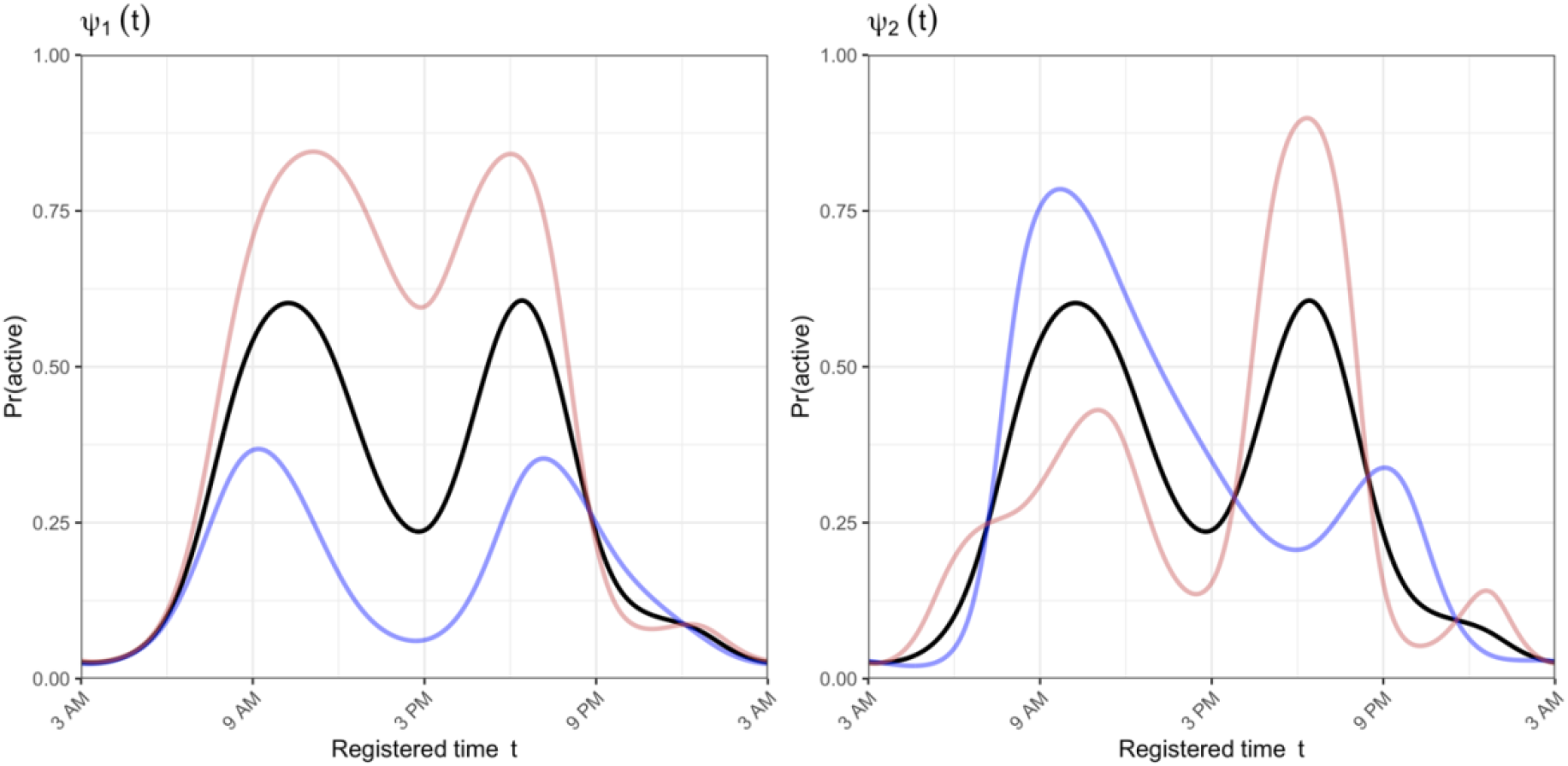
Population mean activity probability profile from registration’s binary FPCA step (black lines), plus or minus some variability (red and blue lines, respectively). The left panel demonstrates the mean +/- 1 standard deviation in the first principal component, and the right panel demonstrates the mean +/- 1 standard deviation in the second component.

The five registration parameters (three warping function slopes and two PC scores) create a five-dimensional parametric space for describing between-subject differences in accelerometry-estimated circadian rhythms, going beyond the continuous one-dimensional morningness-eveningness scale. Extreme values on these five dimensions may be interesting and may yield new, more nuanced description of the complexity of observed chronotypes. For example, extreme values of the warping function slopes classify subjects by their wake and sleep times, leading to chronotypes of “morning lark” vs. “anti-lark” (early vs. late wake time) and “night owl” vs. “anti-owl” (late vs. early sleep time). Extreme values of the first PC score reveal potential chronotypes of “penguin” vs. “hummingbird” (low vs. high overall activity probability); and extreme values of the second PC score reveal potential chronotypes of “rooster” vs. “roadrunner” (most active in the morning vs. afternoon). These potential chronotypes are summarized in Table 1, with corresponding simulated activity probability profiles displayed in Figure 4. In Figure 4, for chronotypes that are defined by warping function slopes, defining features of these chronotypes are most evident before registration, i.e. before the warping functions absorb horizontal variability or differences in sleep and wake times. Conversely, defining features of the PC-score-based chronotypes are more strongly evident after registration, once probability activity profiles have been aligned and all that remains is vertical variability.

**Figure 4.**
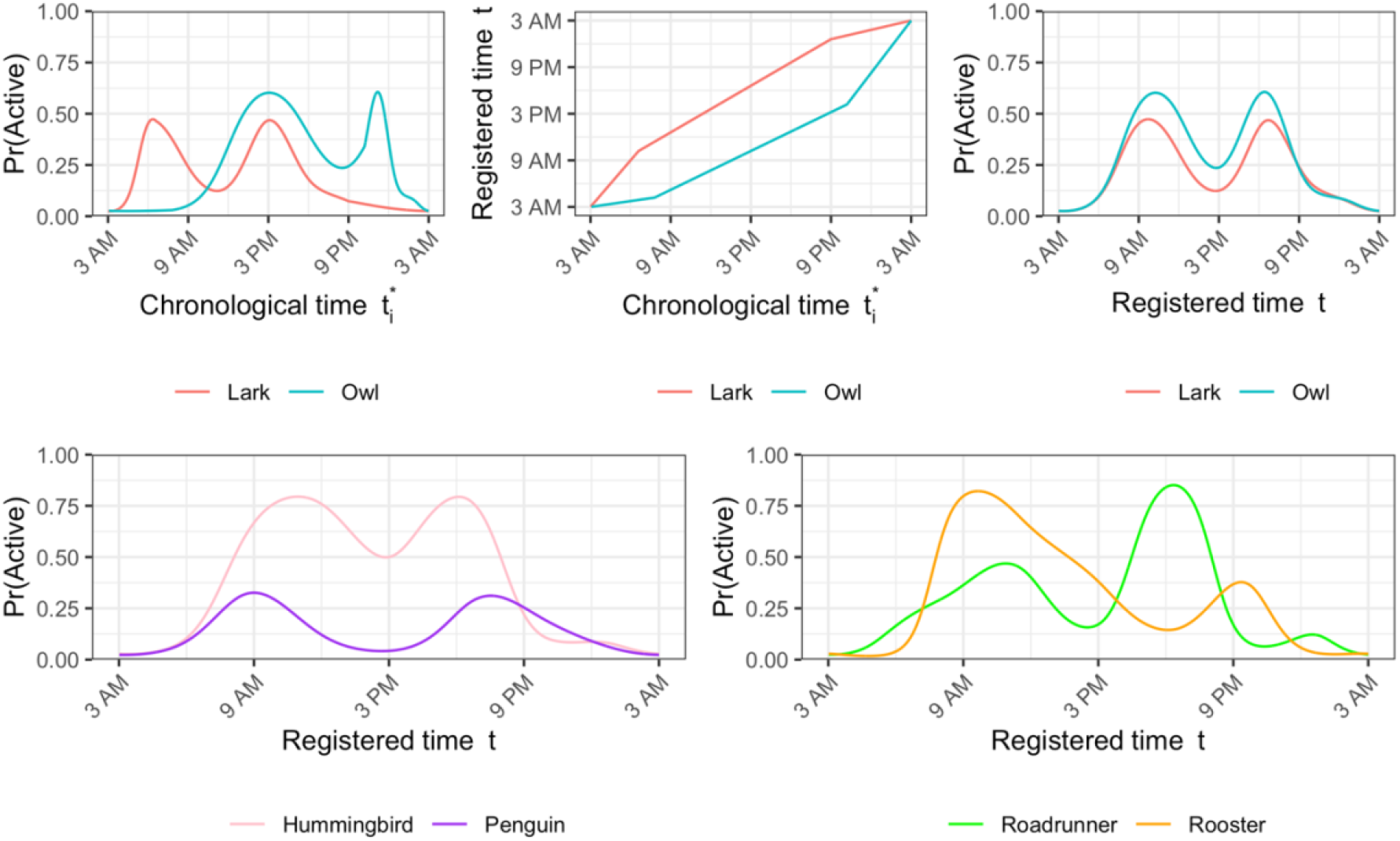
Simulated activity probability profiles from select chronotypes for demonstration. The top row compares a lark and an owl in terms of their profiles in chronological time (top left), warping functions, (top middle), and profiles in registered time (top right). In the bottom left, the profiles of a hummingbird and a penguin are compared in registered time. In the bottom right, the profiles of a roadrunner and a rooster are compared in registered time. These simulated profiles were generated by specifying warping function parameters and PC scores that meet each chronotype’s definition. We refer the reader to Table 1 for these definitions.

**Table 1.** Potential chronotypes defined by cut-off values of registration parameters.

| Registration parameter<br>(for subject $i$) | Values | Proposed<br>chronotype | Defining feature |
| --- | --- | --- | --- |
| $\beta_{1i}$ (1<sup>st</sup> slope of warping function) | High | Lark | Early wake time |
|  | Low | Anti-lark | Late wake time |
| $\beta_{3i}$ (3<sup>rd</sup> slope of warping function) | High | Owl | Late sleep time |
|  | Low | Anti-owl | Early sleep time |
| $c_{1i}$ (1<sup>st</sup> PC score) | High | Penguin | Low overall activity probability |
|  | Low | Hummingbird | High overall activity probability |
| $c_{2i}$ (2<sup>nd</sup> PC score) | High | Rooster | Most active during (registered) morning |
|  | Low | Roadrunner | Most active during (registered) afternoon |

We emphasize that these potential chronotypes are not mutually exclusive, but rather four independent domains on which a subject can present typical or extreme behavior. The ability to define chronotypes across all five (or four, as demonstrated here) dimensions of the registration model may lead to the identification of more sophisticated and distinct subgroups of circadian rhythm patterns compared to existing methods.

## 5. COMPARISON TO EXISTING METHODS

We compared our approach to three existing methods: i) the SPT-window landmark method, ii) the cosinor method, and iii) binary FPCA without registration.

### 5.1. Comparing registration with landmark times

For the SPT-window, we lacked the tri-axial data necessary to calculate arm angle, so we modified the definition of sustained inactivity to be a status of rest for a duration of more than 20 consecutive minutes. This 20minute criterion was chosen post-hoc after visually inspecting its ability to identify approximate sleep and wake times. We estimated two SPT-windows: one during the night between Monday noon and Tuesday noon, and one during the night between Tuesday noon and Wednesday noon. The end of the former SPT-window defined the Tuesday wake time, while the start of the latter SPT-window defined the Tuesday sleep time. Using this approach, we identified wake and sleep times for 424 of the 492 subjects.

Figure 5 contains two panels of lasagna plots (Swihart et al., 2010), in which each row represents a different subject’s rest/activity status over 24 hours, with blue cells representing active minutes and white cells representing rest minutes. In each panel, wake and sleep times for each subject are highlighted in black and the daytime interval midpoint (DIM) for each subject is highlighted in red. Landmark times were first estimated on the chronological time scale (left); and then we evaluated where these landmarks mapped to in registered time (right). Before registration, we first take note that DIMs are not a strong indicator of morningness vs. eveningness; there is considerable overlap in DIM between those who wake later (near the top of the plot) and those who wake earlier (near the bottom of the plot). After registration, we see better alignment in sleep times and wake times as expected, although there is still some between-subject variability, particularly in sleep times. What is more concerning, however, is that the DIM does not consistently map to an intuitively meaningful feature of the registered activity probability profiles. For some patients, the DIM takes place during their midday dip or period of rest. For other patients, it takes place at the height of their morning peak, and still for others, it takes place toward the end of their evening activity peak. The DIM, which is simply calculated as the halfway point between wake and sleep, fails to define any single feature of one’s circadian rhythm.

**Figure 5.**
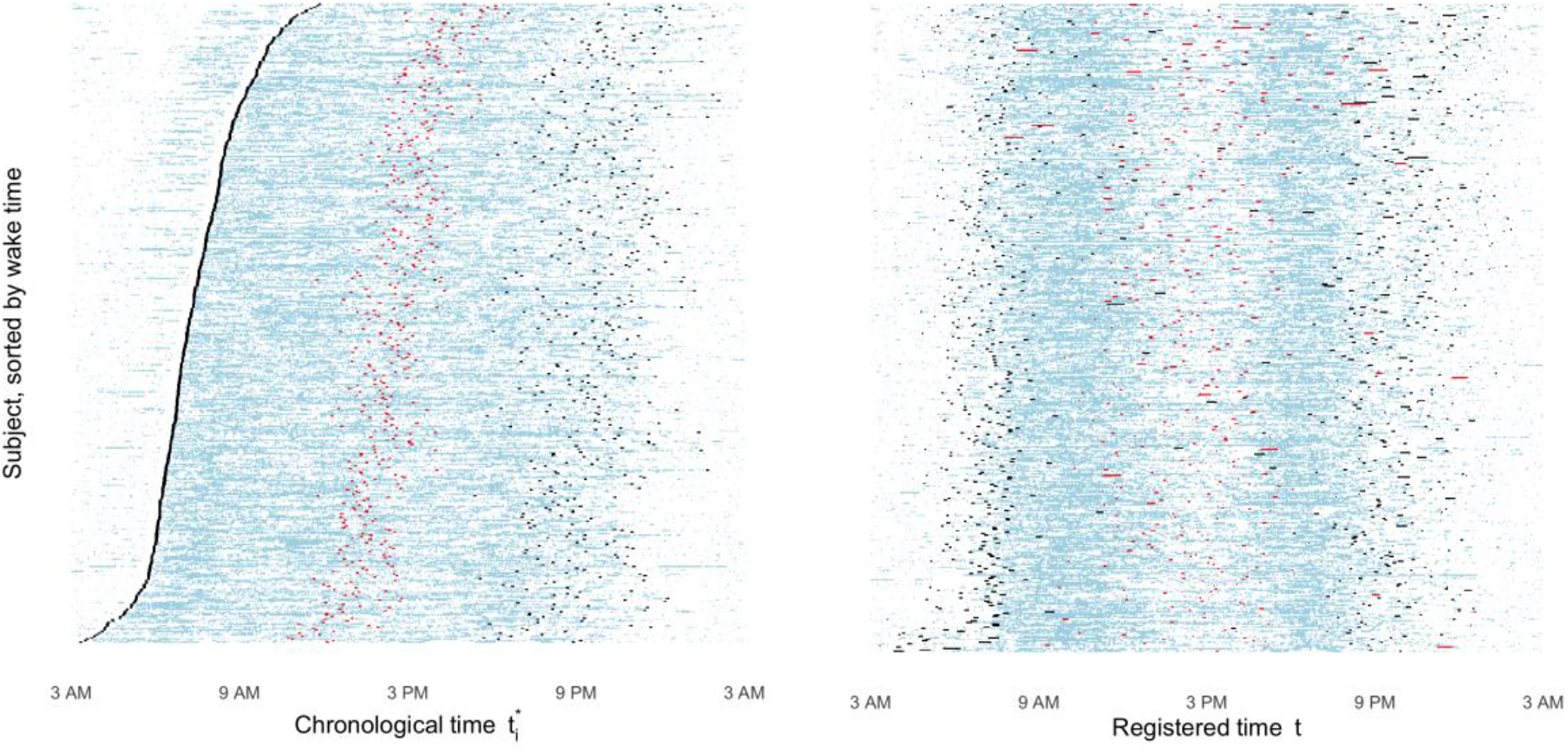
Lasagna plots pre-registration (left) and post-registration (right), with colored markers for landmark times estimated from the (modified) SPT-window approach. In each panel, each row represents a different subject’s binary activity status over time. Blue cells indicate periods of activity, and white cells indicate periods of rest. Sleep and wake times for each subject are marked in black. The daytime interval midpoint (DIM) is marked in red.

### 5.2. Comparing registration with the cosinor model

We fit cosinor curves to each subject’s non-binarized activity count data, noting that the fitted values did not always accurately describe the underlying activity profile, particularly during periods of rest (Supplementary Figure S1). The cosinor models produced three estimated parameters for each subject: the MESOR, the amplitude, and the acrophase. These parameters represent the estimated model-based mean, the peak or maximum deviation from the mean, and the timing of the peak, respectively. Below we examine correlations between these cosinor parameters and the two PC scores of registration’s FPCA step. The top row of Figure 6 shows strong negative correlations between the first PC score and the MESOR and amplitude parameters, which is consistent with expectations. All three cosinor parameters showed at least weak associations with at least one slope parameter as well (Supplementary Figure S2); this highlights that, in contrast to the registration parameters, the cosinor parameters are not designed to explicitly explain one source of variability. In the bottom row of Figure 6, no cosinor parameters are correlated with the second PC, suggesting that this is new information gained from registration that the cosinor model could not uncover, perhaps due to its restricting shape.

**Figure 6.**
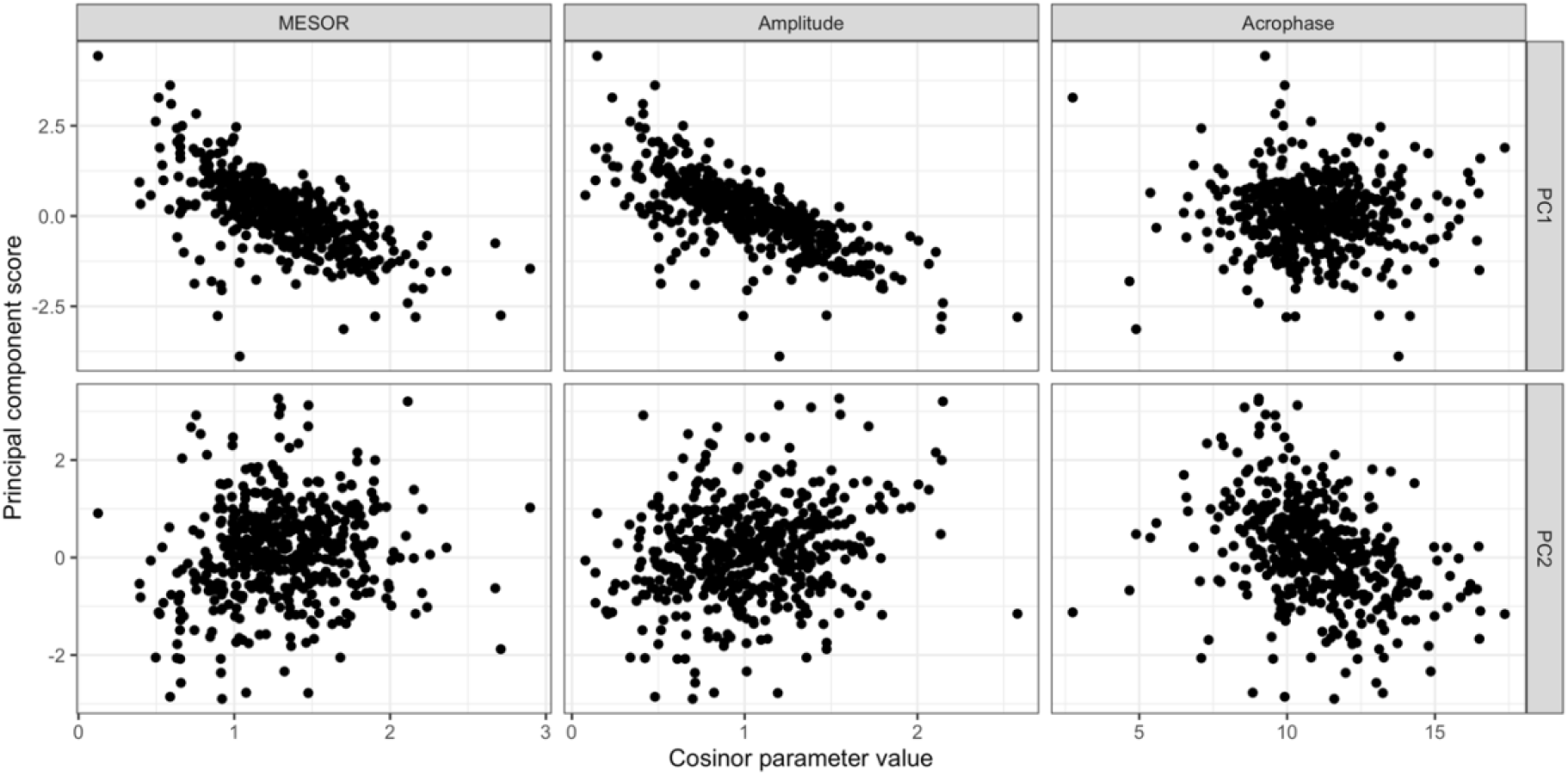
Registration-based principal component scores by cosinor parameter values. Registration-based first and second PC scores by cosinor MESOR (top left and bottom left, respectively); registration-based first and second PC scores by cosinor amplitude (top center and bottom center, respectively); and registration-based first and second PC scores by cosinor acrophase (top right and bottom right, respectively).

### 5.3. Comparing registration with unregistered FPCA

For standalone binary FPCA without registration, we explored a range of different numbers of PCs *K* (from 2 to 4) and determined that *K* = 3 had the best fit and interpretability. The estimated population-level mean is shown in Figure 7 (black lines). In the left panel, colored lines represent the mean plus (blue) or minus (red) one standard deviation in the first PC. Similarly, the middle and right panels show deviations in the second and third PC’s, respectively. The first PC mostly describes the single-dimensional continuum of morningness vs. eveningness, with some additional information about morning activity intensity and the magnitude of midday dip. The second PC describes overall vertical variability in activity probability. The third PC largely describes the timing of the highest peak, but it also contains some remaining information about wake and sleep times that the first PC failed to capture.

**Figure 7.**
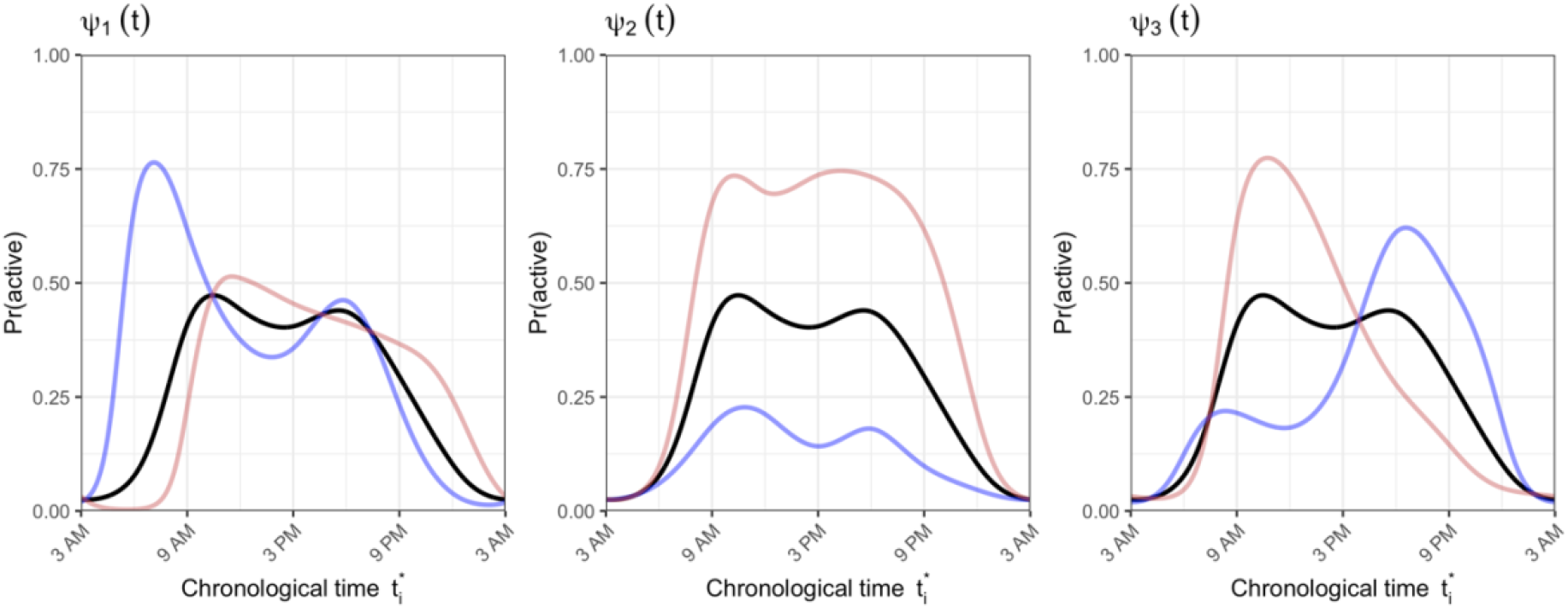
Population mean activity probability profile from binary FPCA without registration (black lines), plus or minus some variability (red and blue lines, respectively). The left, right, and middle panels demonstrates the mean +/- 1 standard deviation in the first, second, and third principal components, respectively.

Binary FPCA without registration is tasked with making sense of all variability, including both horizontal and vertical. Because of this, both the estimated mean and the types of variability explained in these PCs differ from those of registration (Figure 3), which can capture more advanced levels of variability in patterns of activity probability. Binary FPCA without registration captured the general trend of two activity peaks with a midday dip in its population mean activity probability profile, but these features were not as distinct as those revealed with registration. Furthermore, although its resulting PCs can separate subjects with more active mornings versus more active afternoons (similar to the second PC of registration), this information is confounded by claims about the number of peaks, as well as wake and sleep times.

## 6. DISCUSSION

We have demonstrated that registration of 24-hour accelerometric rest-activity profiles can provide new insights into the complexity of chronotypes compared to existing methods by handling distinct sources of variability in daily activity patterns separately. By doing registration, we uncovered limitations in the interpretability of traditional landmark approaches, specifically the DIM estimated from SPT-windows. Further, while we demonstrated an overlap between the results of registration and existing modeling methods such as the cosinor and FPCA, registration revealed novel insights into how a person expends their activity throughout the day.

Though registration provides useful information beyond that obtained from the cosinor method, our goal was to introduce registration as an approach that complements rather than replaces the cosinor method. Like the cosinor method, registration reveals interpretable parameters; but registration provides extra flexibility by allowing the shape of the mean activity probability profile to be driven by the data instead of restricted to a cosine-like shape. That said, we acknowledge that some extensions to the cosinor model, including the multiple-component cosinor (Cornelissen, 2014) and the sigmoidally-transformed cosinor (Marler et al., 2006), also provide flexibility that improves upon the original cosinor approach, albeit in different parametric ways. We aim to introduce registration as another available tool among this set of flexible methods, one which learns the shape of the activity probability profile solely from the data.

When comparing registration to the cosinor method in this analysis, it is important to note the difference in outcomes. The cosinor method modeled the log-transformed activity counts as a continuous outcome, whereas the registration method modeled binary states of active vs. rest. We made this modeling choice because we believe that the binarized activity data is of greater utility than the log-transformed (or untransformed) activity counts for studying circadian rhythms. In part, this is because when using the noisy log-transformed counts, modeling methods may be overly influenced by extreme values which may indicate high levels of activity at a particular point in time, but do not convey information about timing of circadian rhythms. Another limitation of using raw activity counts is the dependence of the results on the definition of activity counts that are often device and manufacturer specific. There may still be some circumstances under which continuous activity count data are preferred, for example in younger populations where scientists may want to understand moderate to vigorous physical activity (MVPA). For these research questions, methods for registering Gaussian data have been developed and are available in the “registr’ package in R (Wrobel et al., 2020). However, for the reasons listed above, we believe binarized rest-activity profiles are more useful for studying circadian rhythms. Our approach to understanding circadian rhythms using binarized accelerometer data is compelling and novel, particularly the insights gained from the second PC obtained with our method.

While registration may seem vastly different from methods commonly used in chronobiology research such as the cosinor method and FPCA, we see many parallels between them. Obviously FPCA and registration are related in the sense that FPCA itself is a step in the registration method. However, a more fundamental similarity across all three methods is the treatment of accelerometer data as functional data across the 24-hour time span. The equations behind the registration, cosinor, and FPCA models are all written as functions of time. From this perspective, we see that the concept of registration is not so far-fetched from the types of methods that are already commonly applied in chronobiology. Finally, our approach is unique in that it borrows information across all subjects. Thus, the approach is potentially more powerful compared to those that focus only on a subject-specific data.

One limitation of this method, as well as the cosinor method and FPCA, is that they require a full 24 hours of activity data. For older studies this data may not be available since subjects have sometimes been instructed to remove the accelerometer device during sleeping hours (Troiano et al., 2014; Urbanek et al., 2018). However, 24-hour data collection of wrist-worn accelerometer data is becoming increasingly more popular in the physical activity and sleep research communities (Doherty et al., 2017; van Hees et al., 2018). Access to full 24-hour activity profiles will be less of a challenge looking forward.

A second limitation, specific to registration, is the assumption that the method is only useful if the population has an underlying mean that is shared, at least to some extent, across subjects. In our analysis, the mean suggested a morning activity peak, and midday dip, and an afternoon activity peak, with rest during nighttime hours. The structure we assume is reasonable and supported by the BLSA data in the sense that many warping functions lie close to the identity line, suggesting minimal warping required to align these subjects. Furthermore, this biphasic structure has also been identified as an FPCA component of other large-scale accelerometer studies such as NHANES (Leroux et al., 2019) and the Osteoporotic Fractures in Men (MrOS) study (Zeitzer et al., 2018) as well as in monkeys (Leroux, 2015). It is also important to note that the interpretations of our registration results, i.e. the potential chronotypes presented in Table 1 and Figure 4, may be specific to this analysis. Performing registration on other datasets or with other analysis specifications, such as the choice of starting point for the 24-hour period, may yield PCs with different interpretations. Here we have defined the daily cycle to start at 3AM; we would expect a different mean function with different PCs and warping functions if the analysis were re-run using a different starting time. Future work is needed to validate our potential new chronotypes using other data sources.

As a final limitation, we highlight that each of these analysis approaches utilize only a single day of data per subject. To make robust claims about one’s chronotype, we would ideally analyze several days of accelerometer data and estimate within-subject day-to-day stability, variability, and regularity of rest-activity rhythms, topics that are currently under active investigation (Phillips et al., 2017). Future work is needed to create longitudinal extensions of the registration method.

Future work is also warranted to explore subgroups of people based on their circadian rhythms as determined by their subject-specific registration parameters. In chronobiology, terms such as “larks” and “owls” are commonly used to describe morning and evening chronotypes (Roenneberg et al., 2003). Furthermore, these chronotypes are associated with many health outcomes (Partonen, 2015). We have shown that registration parameters can distinguish these and additional chronotypes, describing more nuanced features of one’s daily activity patterns beyond the timing of falling asleep and waking. Furthermore, these chronotypes can be defined on a 5-dimensional continuum, similar to but more flexible than the well-known single-dimensional morningness-eveningness chronotype continuum (Partonen, 2015). Future work is needed to both validate these dimensions using independent data sources and to explore associations between these potential chronotypes and health outcomes.

