## Supplementary for "Scale-invariant time registration of 24-hour accelerometric rest-activity profiles and its application to human chronotypes"

Supplementary material

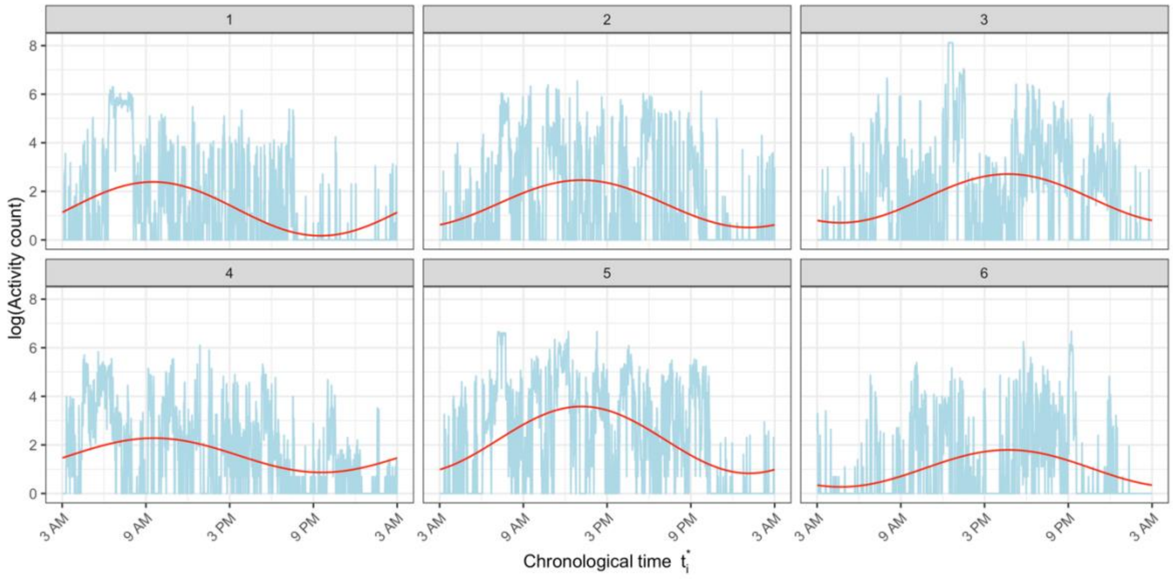


*Figure S1. Fitted cosinor curves (in red) overlayed on log-transformed activity count data (in blue) for 6 select subjects from the BLSA dataset.*

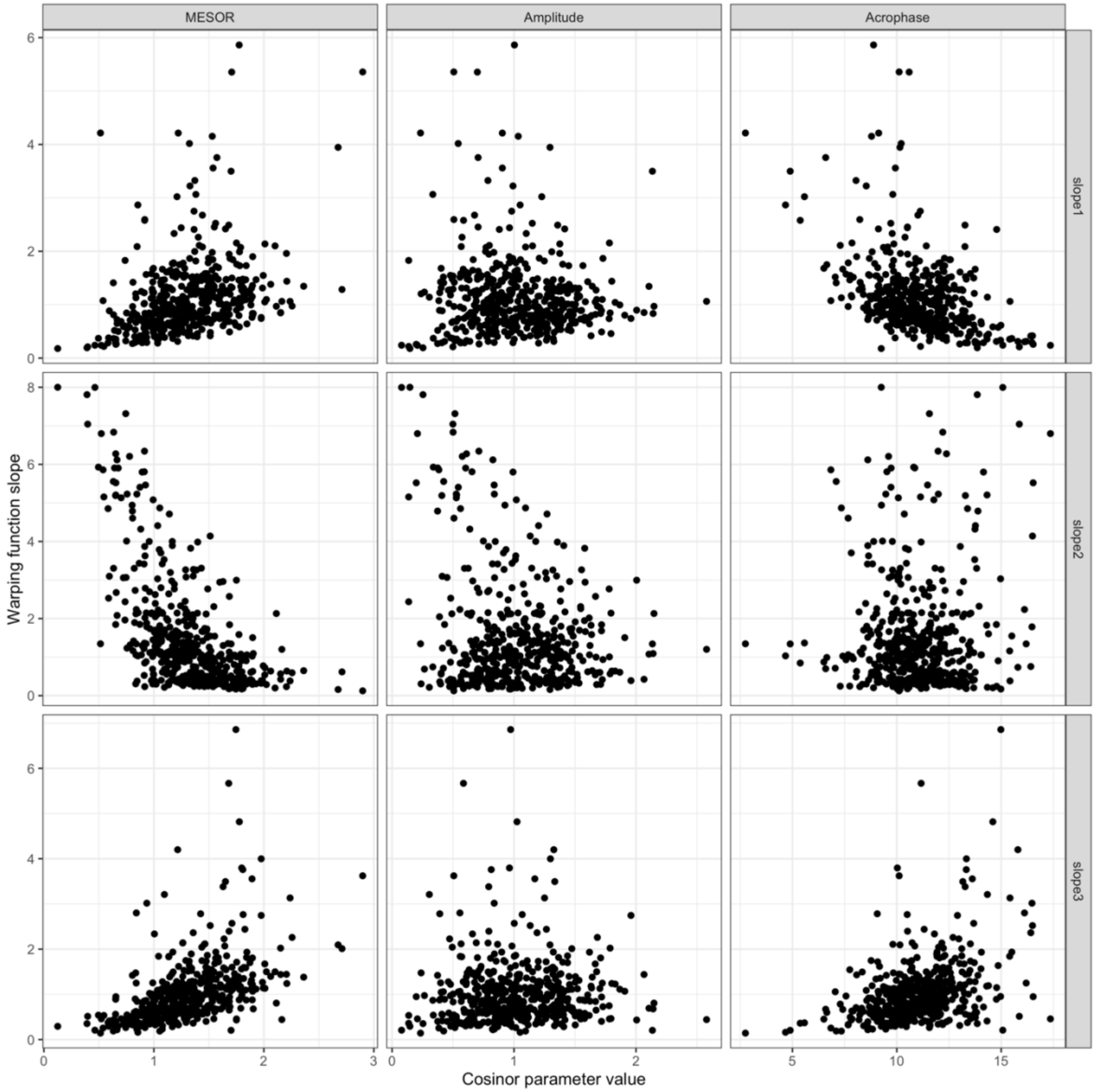


*Figure S2. Registration warping function slopes by cosinor parameter values. Registration first, second, and third slopes by cosinor MESOR (top left, middle left, and bottom left, respectively); registration first, second, and third slopes by cosinor amplitude (top center, middle center, and bottom center, respectively); and registration first, second, and third slopes by cosinor acrophase (top right, middle right, and bottom right, respectively).*
